# Mutation and ACE2-induced Allosteric Network Rewiring in Delta and Omicron SARS-CoV-2 Spike Proteins

**DOI:** 10.1101/2025.11.12.688084

**Authors:** Mandira Dutta, Gregory A. Voth

**Affiliations:** Department of Chemistry, Chicago Center for Theoretical Chemistry, Institute for Biophysical Dynamics, and James Franck Institute, The University of Chicago, Chicago, IL 60637

## Abstract

The spike protein of severe acute respiratory syndrome coronavirus 2 (SARS-CoV-2) mediates viral entry by binding its receptor-binding domain (RBD) to the host receptor ACE2. Spike mutations in different variants have been experimentally shown to influence the rate of conformational transitions and alter viral infectivity. In parallel, both experimental and computational studies have reported the presence of long-range allosteric communication within the spike protein, suggesting that such mutations may also affect allosteric signaling pathways involved in viral function. A detailed understanding of the allosteric residue network is essential for rational antiviral drug design. In this study, we performed extensive atomistic molecular dynamics (MD) simulations of the spike proteins from the Delta and Omicron variants in both ACE2-bound and unbound states. By integrating linear mutual information (LMI) calculations and graph theory-based analysis, we delineated the long-range allosteric communication networks embedded within the spike protein. Betweenness centrality metrics enabled the identification of residues that act as key mediators of information flow. Notably, ACE2 binding markedly enhances allosteric coupling throughout the spike. We identified three key linkers, Link1 (NTD-RBD), Link2 (RBD-SD1), and Link3 (SD2-FP), as primary mediators of allosteric communication. Delta exhibits stronger signaling through Link1 and Link2, whereas Omicron redirects communication via Link3. While Delta maintains localized connectivity within the S1 domain but loses long-range contact with the S2 core, Omicron forms a broader yet weaker S1 network and establishes long-range coupling. We propose that the N856K and T547K mutations reshape the conformational landscape, reconfiguring allosteric communication pathways in Omicron. Furthermore, our analysis reveals distinct domain-level allosteric couplings in Delta and Omicron, pointing to variant-specific differences in fusogenicity and immune evasion. By mapping key allosteric sites and mutation-induced conformational shifts, our study may provide a framework for developing robust antiviral strategies resilient to future emerging SARS-CoV-2 variants.

**Significance:** Receptor engagement at the RBD rewires long range allostery in the SARS CoV 2 spike. Using LMI and graph theory-based analyses, we map communication paths and pinpoint residues that govern spike opening and infection. ACE2 binding globally strengthens coupling, but energy and signals propagate along variant specific routes: Delta biases toward openness, channeling binding energy into RBD opening; Omicron remains less open, routing energy to the S2 core to prime fusion. Information flows through three linkers - Link1 (NTD to RBD), Link2 (RBD to SD1), and Link3 (SD2 to FP), with Delta emphasizing Link1/Link2 and Omicron shifting to Link3 and CD connections. We suggest that the N856K and T547K mutations reshape this landscape in Omicron.

## Introduction

Allostery is a fundamental process by which protein machines sense and respond to environmental perturbations, such as ligand binding, mutations, or posttranslational adaptations, and relay these signals to distant sites, eliciting functional responses.(1–5) The spike (S) protein of severe acute respiratory syndrome coronavirus 2 (SARS-CoV-2), the primary target for neutralizing antibodies(6) and the focus of this study, exemplifies this principle. As a large and complex biomolecular machine, the S protein undergoes extensive conformational changes to mediate fusion of the viral and host membranes, triggered by host receptor attachment at its receptor binding domain (RBD).(7,8) The communication between the RBD and distant subunits regulates the protein’s conformational dynamics, highlighting the presence of allosteric signaling within the S protein.(9)

Since its emergence, the SARS-CoV-2 virus has had a profound global impact, resulting in over 800 million reported cases and approximately 7 million fatalities worldwide (https://ourworldindata.org/coronavirus). Although vaccination campaigns have significantly reduced disease severity and mortality, the rapid evolution of SARS-CoV-2 variants, including Alpha, Beta, Gamma, Delta, and an expanding range of Omicron sublineages such as BA.1 to BA.5, BQ, XBB, EG.5, and BA.2.86, continues to challenge vaccine efficacy (https://www.ecdc.europa.eu/en/covid-19/variants-concern). While numerous studies have focused on mutations within the RBD and their impact on the interaction of the RBD with the host cell receptor ACE2,(10,11) mutations in the RBD and other regions of the spike protein may also influence distal subunits, thereby altering the overall conformational dynamics of the S protein.(12–14) Despite significant advances, our understanding of allosteric regulation remains limited, with the underlying molecular mechanisms largely unresolved, primarily due to the structural complexity of the S protein. This knowledge gap continues to hinder the rational design of allosteric targeted therapeutics.

The virus consists of a genomic RNA within a protective protein shell, composed of four major structural proteins: Spike (S), Envelope (E), Membrane (M), and Nucleocapsid (N) proteins.(15–18) The S protein is a heavily glycosylated trimeric membrane protein that anchors the viral membrane with its transmembrane domains(19–21) (Figure 1a). In its prefusion state, the spike glycoprotein comprises two subunits: S1, responsible for receptor binding, and S2, which mediates viral host membrane fusion (Figure 1a). The S1 subunit of the spike protein consists of the N-terminal domain (NTD), the receptor binding domain (RBD), and subdomains 1 and 2 (SD1/2), which act as hinge regions modulating RBD movement. The S2 subunit contains the fusion peptide (FP), two heptad repeats (HR1 and HR2), the transmembrane domain (TM), and the cytoplasmic domain (CD)(19,20) (Figure 1b). Viral entry is initiated through a stepwise process involving conformational transitions of the RBD from the “down” to the “up” state upon ACE2 binding, followed by cleavage at the S1/S2 furin site, leading to S1 shedding, and subsequently, cleavage at the S2’ site.(22,23) A linker between the NTD and RBD (residues 293-329), referred to here as Link1, was found to correlate with RBD opening.(24–26) In a computational study by Kearns *et al*., (27) this linker was denoted as N2R. They also identified another linker (residues 522-536), which connects the RBD and NTD via a disulfide bond and was named R2N; we refer to this as Link2. They found that both linkers play important roles during RBD opening. Additionally, we identify a third linker between SD2 and FP (residues 686-715) that connects the S1 and S2 subunits through a flexible disordered segment, which we designate as Link3. We hypothesize that all three linkers are important for allosteric communication within the spike protein. A previous coarse-grained molecular dynamics (CG MD) study (28) has further shown that shedding of the S1 domain and activation of the S2 core for viral fusion depends on the cooperative binding of multiple ACE2 molecules to the same S protein trimer. After the removal of S1 from the upside-down pyramid-like prefusion S, S2 adopts an elongated rod-like structure, where the HR1 and CH helices form a continuous extended helix from a previously “U”-shaped bent structure, yielding a rigid six-helix bundle.(29,30) A recent study examined the conformational changes in the SARS-CoV-2 S2 trimer and designed a stable, prefusion-locked S2-based immunogen.(31)

**Figure 1.**
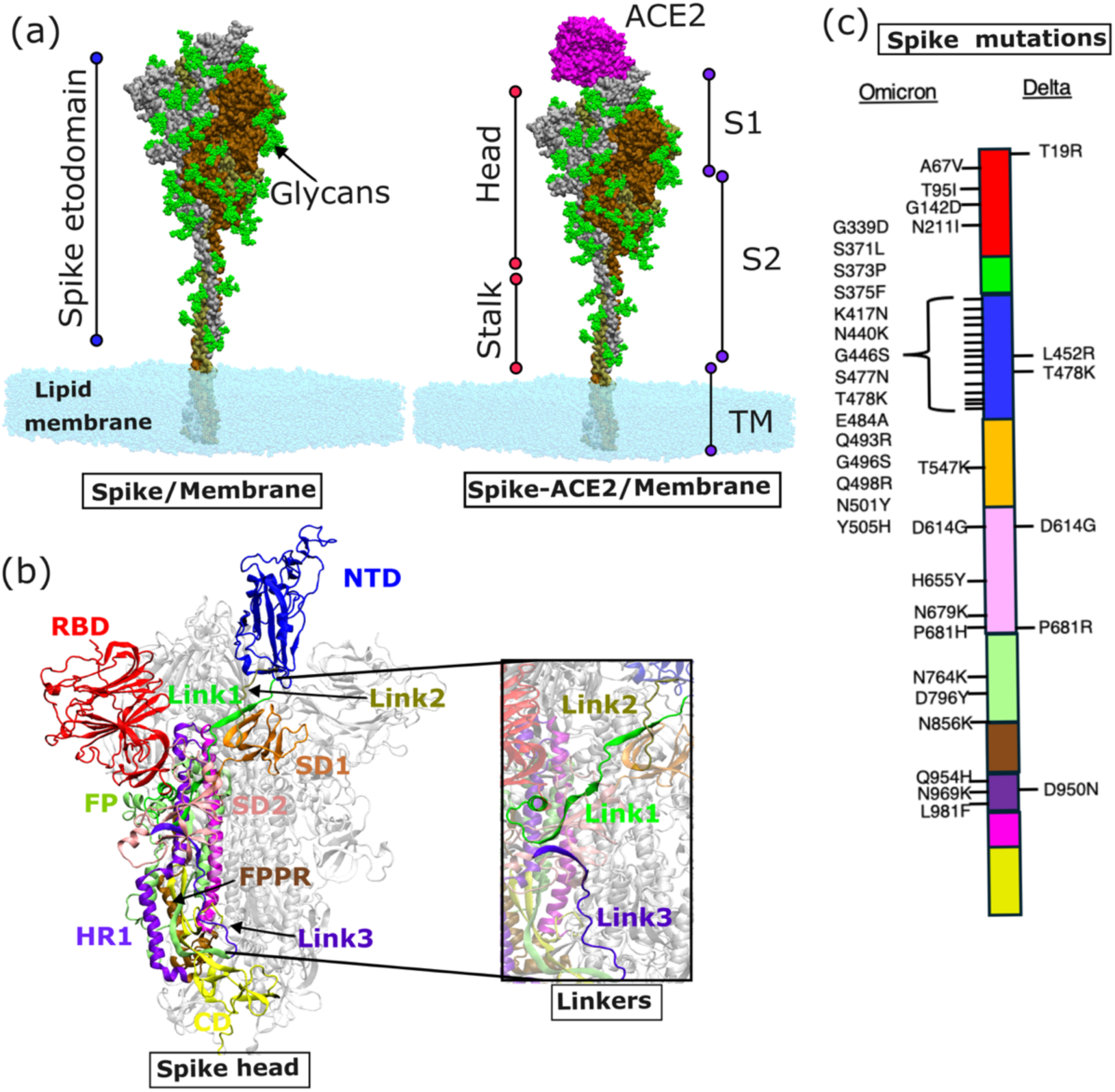
Structural Overview of the SARS-CoV-2 Spike Protein and Variant-Specific Mutations. (a) Structure of the SARS-CoV-2 spike protein in ACE2-unbound and ACE2-bound states, embedded in a lipid membrane. The S1 domain spans residues 14-685, the S2 domain spans residues 686-1147, and the stalk, including the TM region, spans residues 1148-1270. (b) Different subdomains of the spike protein are shown in distinct colors. S1 consists of the NTD (residues 14-292), Link1 (residues 293-329), RBD (residues 330-521), Link2 (residues 522-536), SD1 (residues 537-590), and SD2 (residues 591-685). The S2 domain comprises Link3 (residues 686-715), FP (residues 716-854), FPPR (residues 855-911), HR1 (residues 912-984), CH (residues 985-1034), and CD (residues 1035-1147). (c) Mutations in the Delta and Omicron variants mapped onto their respective subdomains.

Mutations have significantly influenced the conformational dynamics of SARS-CoV-2 variants. Studies have shown that the initial D614G spike loses the stabilizing interaction between D614 (S1) and K854 (S2), which affects the FPPR loop and the furin cleavage site, thereby contributing to the increased stability of the prefusion state in the D614G structure.(32,33) Furthermore, a population shift toward the one RBD-up open conformation was observed in the D614G variant, providing a potential explanation for its enhanced infectivity.(33)

The Delta variant carries two mutations in the RBD and two in the NTD, with a notable mutation at position 681(34) (Figure 1c). The P681R mutation in Delta has been associated with improved furin-mediated cleavage of the S protein.(35,36) While *in vitro* assays indicate that this mutation does not directly enhance the infectiousness of the Delta variant, *in vivo* studies in hamsters suggest that it contributes to increased pathogenicity compared to the wild-type SARS-CoV-2 strain.(36,37) In contrast, Omicron mutations are densely concentrated in the NTD and RBD regions, along with additional mutations in the RBD hinge regions and S2 domain(38) (Figure 1c). Omicron variants have also exhibited substantial resistance to spike-targeting monoclonal antibodies, reducing the efficacy of previously authorized therapeutic treatments for SARS-CoV-2 infection.(39) The Omicron S protein exhibits slightly reduced S1/S2 cleavage, indicating that the N679K and P681H mutations at the furin cleavage site do not enhance S protein processing.(40,41) Additionally, five mutations (T547K, H655Y, N856K, Q954H, and N969K) contribute to decreased fusogenicity, while the L981F mutation appears to counterbalance this effect by partially restoring fusogenicity.(42)

Both experimental and computational studies have reported the presence of allosteric communication in the spike protein.(9,13,26,43) A previous study explored the dynamics of the free S protein and the S:ACE2 complex to understand the allosteric effects of ACE2 binding on distal stalk regions and protease docking sites.(13) ACE2 engagement initiates the generation of S2′ fragments in target cells, a crucial proteolytic step that facilitates spike-mediated membrane fusion. Qiao and Olvera de la Cruz investigated how mutations and polybasic cleavage sites, located ~10 nm away from the RBD, influence the RBD-ACE2 interaction.(44) Chen et al. studied ACE2-induced allosteric activation in SARS-CoV and SARS-CoV-2 spike proteins using hydrogen/deuterium-exchange mass spectrometry, revealing that ACE2 binding triggers allosteric changes from the RBD to cleavage sites and the fusion peptide.(45) Thirumalai and co-workers reported that ACE2’s substrate-binding cleft, located far from the interface, opens when the spike-ACE2 complex dissociates.(46) They also found that RBD binding enhances movements in this cleft, suggesting a connection between SARS-CoV-2 binding and ACE2’s enzymatic activity. Other studies have shown how human antibodies induce conformational and allosteric changes in the spike protein, revealing distinctions between weak, moderate, and strong neutralizers.(47,48) Ray et al. identified allosteric sites in the spike protein by analyzing long-range correlations between the RBD and distant residues.(26) Using allosteric signaling networks within the spike protein, another study identified potential variant-driving mutations and novel distal sites that could be targeted by allosteric drugs.(43) Dokainish et al. revealed key inter-domain interactions that drive transitions between Down, one-Up, one-Open, and two-Up-like states, highlighting cryptic pockets and functionally relevant intermediate states.(49) A more recent study demonstrated how SARS-CoV-2 spike mutations, including D614G, influence RBD opening by reshaping the conformational landscape of the S protein.(27) While numerous studies have recognized allosteric regulation in the spike protein, the detailed mechanism remains unclear. In particular, how mutations in SARS-CoV-2 variants alter the pathways that transmit signals from the RBD to distal allosteric sites upon ACE2 binding has yet to be fully elucidated.

To address this gap, the present study investigates how mutations in the Delta and Omicron BA.1 (hereafter referred to as Omicron) variants alter allosteric communication upon ACE2 binding within a spike-membrane system. We performed extensive atomistic MD simulations of the Delta and Omicron spike proteins embedded in a lipid bilayer, in both ACE2-bound (Delta_+ACE2_ and Omicron_+ACE2_) and unbound (Delta_−ACE2_ and Omicron_−ACE2_) states (Figure 1a). We characterized the distinct conformational selections upon ACE2 binding in the Delta and Omicron RBDs based on two selected collective variables (CVs). To identify key residues, we constructed a protein connectivity network using graph-theoretical analysis based on linear mutual information (LMI) metrics derived from MD trajectories. Betweenness centrality (BC) was used to identify residues that transmit the maximum allosteric signal within the spike protein. Our analysis reveals that ACE2 binding significantly enhances allosteric coupling in the spike. We also observed that Link1, Link2, and Link3 play major roles in transmitting allosteric communication within the S1 domain and between the S1 and S2 domains. We propose that the Omicron-specific mutations N856K and T547K reshape the allosteric pathways in the spike. Furthermore, our results reveal the gain and loss of domain-wise allosteric coupling, particularly in the linker regions, highlighting distinct signaling routes in Delta and Omicron.

We hypothesize that Link1 regulates RBD opening and epitope gating, Link2 clamps RBD-SD1 hinge to govern S1 retention versus shedding, and Link3 communicates ACE2 binding to the S2 core to regulate priming/fusion. These mechanistic roles align with variant-dependent spike behaviors, where Delta shows stronger coupling through Link1 and Link2 to favor RBD opening and enhanced S1/S2 cleavage,(36) while Omicron reinforces Link3-mediated communication that supports a less open trimer and stronger S1/S2 coupling.(24) By pinpointing key allosteric sites and mutation-induced conformational shifts, our findings provide a structural and energetic framework that connects allosteric communication to functional diversity across variants and may inform the design of antiviral strategies resilient to future evolutionary changes.

## Methods

### System Preparation

The SARS-CoV-2 spike protein is a fully glycosylated, large, and complex homotrimer composed of three identical monomers. Its ectodomain, shaped like an upside-down pyramid (residues 14-1162), is connected to a long stalk region (residues 1163-1273) that is embedded in the viral membrane(8) (Figure 1). The spike protein exists in two major conformational states: open and closed. In the open state, one, two, or all three RBDs adopt the ‘up’ conformation, whereas in the closed state, all RBDs remain in the ‘down’ conformation.(22,23) In the current study, we considered the one RBD-up conformation as the open state of the spike. The one RBD-up structures for Delta and Omicron were taken from PDB IDs 7SBL(34) and 7TO4(29), respectively. Figure 1c summarizes the mutations across different regions of the S protein. To model the spike trimer in complex with the ACE2 dimer, we used the crystal structures of the RBD/ACE2 complexes from PDB ID 8HRK(50) for Delta and PDB ID 7WPB(51) for Omicron. The RBD-up domain of each spike structure was aligned with the RBD domain of the RBD-ACE2 complex to generate the ACE2-bound spike conformations. Both spike PDBs (7SBL and 7TO4) lacked the stalk domains; to model the complete spike ectodomain and stalk, we incorporated the stalk region from the structural model developed by Casalino et al.(21) Missing residues in the spike protein structures were specified using Modeller 9.19.(52)

A recent study by Shajahan et al. compared the site-specific glycosylation profiles of the wild type and five SARS-CoV-2 variants (Alpha, Beta, Gamma, Delta, and Omicron).(53) Although both N- and O-glycosylation sites are highly conserved, especially in the RBD region, they observed notable differences in the glycosylation patterns across variants, with Omicron exhibiting the highest deviation. Specifically, Omicron was found to have fewer oligomannose-type glycans compared to Delta. Based on this study, we modeled the glycosylation profiles of the Delta and Omicron spike proteins accordingly. Each Delta monomer contains 20 N-glycosylation sites and one O-glycosylation site, whereas each Omicron monomer has 18 N-glycosylation sites and one O-glycosylation site. To construct the fully glycosylated spike proteins, we used the Glycan Reader & Modeler tool in CHARMM-GUI.(54)

For membrane modeling, we constructed a symmetric bilayer lipid membrane patch of 25 × 25 nm² with the composition POPC (45%), POPE (20%), Cholesterol (15%), POPI (10%), and POPS (10%). This composition was chosen to mimic a physiologically relevant mixed bilayer representative of the endoplasmic reticulum–Golgi intermediate compartment (ERGIC).(21) The S/lipid membrane system was generated using the Membrane Builder in CHARMM-GUI.(55) A fully hydrated lipid bilayer was constructed around the S protein using TIP3P water molecules, positioning the transmembrane domain near the bilayer midplane. System neutrality was maintained by adding 0.15 M KCl.

In this study, we prepared four S/lipid membrane systems: Delta-S, Delta-S+ACE2, Omicron-S, and Omicron-S+ACE2. Each system contains approximately 2.4 million atoms in the absence of ACE2 and around 3 million atoms in the presence of ACE2.

### MD Simulations

We carried out atomistic MD simulations using GROMACS with the CHARMM36m force field.(56,57) The system was energy-minimized and equilibrated through the six-step protocol provided by the CHARMM-GUI.(55) Simulations were performed with a 2fs integration time step under periodic boundary conditions. The temperature was held constant at 310.15 K using a Nosé–Hoover thermostat(58,59) with a 1.0 ps coupling time. During the initial relaxation, pressure control was maintained with the Berendsen barostat(60), followed by production runs using the Parrinello-Rahman barostat(61,62) applied semi-isotropically with a compressibility of 4.5 × 10⁻⁵ bar⁻¹ and a coupling constant of 5.0 ps. Non-bonded interactions were smoothed between 1.0 and 1.2 nm, and long-range electrostatics were computed using the Particle Mesh Ewald method(63). Hydrogen bond lengths were constrained using the LINCS algorithm.(64) We completed 1.6 µs of production simulation on Frontera at the Texas Advanced Computing Center (TACC) and Midway2 (University of Chicago RCC) using GROMACS-2019. We simulated three replicas for each system.

### Analysis

#### Linear Mutual Information Analysis

In information theory(65), entropy is a fundamental measure that quantifies the uncertainty associated with a discrete random variable *X*. The variable *X* can take outcomes *x*_1_, *x_2_,* …, *x_n_,* with probabilities defined by the mass function *p*(*x*). When information about another random variable *Y* is available, the conditional entropy of *X* can be evaluated. The reduction in uncertainty about *X* due to knowledge of *Y* is defined as mutual information (MI), which quantifies the degree of statistical dependence between the two variables, as described by equation 1.

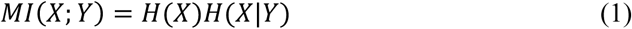

Here, *H*(*X*) is the entropy of *X*, and *H*(*X*|*Y*) is the conditional entropy given *Y*. Mutual information drops to zero when *X* and *Y* are independent, meaning knowledge of one does not reduce uncertainty about the other. As mutual information is symmetric, this result applies equally to both variables.

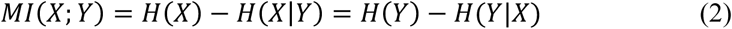

MI in terms of joint entropy is expressed as:

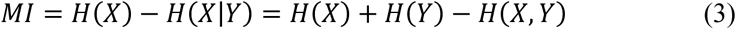

The decrease in entropy described in Eq. (1) can alternatively be expressed in terms of relative entropy, which quantifies the divergence between the joint probability distribution *p*(*x, y*) and the product of the marginal distributions *p*(*x*)*p*(*y*). This interpretation highlights mutual information as a measure of how far the true joint distribution deviates from what it would be under independence. The mutual information is defined as:

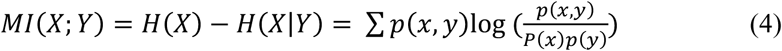

The mutual information can be reformulated as a specific instance of relative entropy. This formulation has been widely applied to identify and quantify correlated motions in MD trajectories. In this context, the variables *x* and *y* in Eq. (4) are replaced by displacement vectors of Cα atoms corresponding to residues i and j, defined as *x_i_* = *r_i_* − 〈*r_i_*〉. The discrete probability mass functions are replaced by continuous probability density functions, and the summation is substituted with integration.

Assuming quasi-harmonic behavior of atomic fluctuations, a Gaussian approximation is applied to both the joint and marginal probability density functions of atomic displacements. Under this assumption, the mutual information, representing the reduction in entropy, can be computed analytically, leading to the formulation of LMI as given in Eq. 5:

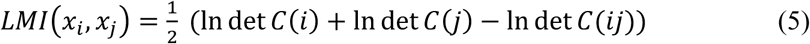

Where *C*(*ij*) = 〈(*x_i_*, *x_j_*)*^T^* (*x_i_*, *x_j_*)〉, and 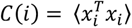

To quantify the strength of correlation in a normalized manner, the LMI-based correlation coefficient is defined as:

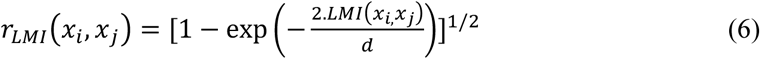

Where *d* = 3 accounts for the dimensionality of the atomic displacements. The normalized *r_LMI_* ranges from 0 (no correlation) to 1 (perfect correlation).

We used the MDAnalysis Python package to process the MD trajectories and extract the Cα atom displacement vectors of each residue. By employing NumPy and SciPy for covariance and log-determinant evaluations, we computed LMI using a custom implementation based on the Gaussian approximation.

### Betweenness Centrality Calculations

Betweenness centrality is a graph-theoretical metric that quantifies the extent to which a node or edge facilitates information flow within a network.(66) Specifically, if a node serves as a bridge along the shortest path between two other nodes, its BC value reflects the frequency with which it lies on such paths.

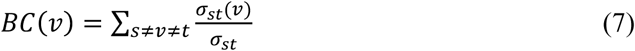

where *σ_st_* denotes the total number of shortest paths between nodes *s* and *t,* and *σ_st_*(ν) is the number of those paths that pass through the node *v*. This measure captures how often node *v* acts as a bridge along the shortest paths connecting other nodes in the network. We used the NetworkX Python package for BC calculations.(67)

Principal Component Analysis (PCA), contact analysis, and residence time calculations were performed using GROMACS modules(56) and the MDAnalysis package.(68) For salt bridge contacts, we considered charged residues, ARG, LYS, ASP, and GLU, with a cutoff distance of 0.4 nm. VMD and ChimeraX were used for visualization purposes.(69,70)

## Results

In our study, we built a full-length glycosylated SARS-CoV-2 spike for Delta and Omicron variants of one RBD up conformation in both bound and unbound states of ACE2. We used the Cryo-EM structures of PDB IDs 7SBL(34) and 7TO4(38) for the Delta and Omicron, respectively. To model the S/ACE2 complex, we used the RBD/ACE2 complex crystal structure of PDB ID: 8HRK for Delta(50), and PDB ID: 7WPB for Omicron (51). We performed 1.6 μs atomistic MD simulations of the trimeric spike protein of Delta, and Omicron variants with the receptor ACE2 bound and unbound states (Figure 1a). The spike protein is fully glycosylated and embedded into the symmetric lipid bilayer membrane with composition POPC (45%), POPE (20%), Cholesterol (15%), POPI (10%), and POPS (10%). We simulated three replicas for each system. The details of the structure preparation and simulations are provided in the *Method* section.

### Modulation of conformational landscape upon ACE2 binding in the spike variants

The structural adaptability of spike proteins is key to their ability to engage host receptors. In the prefusion state, the spike protein primarily transitions between two dominant conformations: a closed state, in which all the RBDs are buried, and an open state, in which one or more RBDs are exposed.(22) These canonical states are connected via an effectively continuum of intermediate conformations. For the open state simulations, we have used one RBD-up (chain A) and two RBD-down protomers (chain B and chain C) of the spike. To characterize the different conformational selection of these variants upon receptor binding, we have defined two collective variables, (1) PC1, the first principal component calculated using the Cα atoms of the S1 domains, is used to describe the collective motion of the S1 domain of the spike protein. Figure S1 (Supporting Information) shows the projections onto PC1 and PC2. Notably, PC1 captures a substantial fraction of the total conformational variance, indicating that the global spike dynamics can be effectively represented by a single collective coordinate. Figure S2 illustrates the PC1 motions in different spike systems. (2) The distance between residue TYR449 in the RBD and residue LYS986 from the tip of the HR1 domain (S2 tip), reflecting the extent of RBD opening (Figure 2a). We observed that the two spike variants, in presence and absence of ACE2, traverse different phase space as defined by our two CV coordinates. Based on the extent of RBD opening across the four systems, which ranges from 65 A° to 80 A°, we defined three distinct states of opening. A distance value of ~65 Å is defined as the ‘open’ state, ~75 Å as the ‘full-open’ state, and any further opening beyond ~75 Å as the ‘extra-open’ state. Interestingly Delta_+ACE2_ displays the maximum RBD opening with an extra-open state and a narrow PC1 range (Figure 2b), indicating a more expanded and stable RBD opening upon ACE2 binding. Surprisingly Delta_−ACE2_ and Omicron_+ACE2_ sample states with the least opening of RBDs. Omicron_−ACE2_ exhibits an intermediate opening of RBD with the full-open state (Figure 2b). Delta_−ACE2_ exhibits broader PC1 distribution reflecting greater conformational heterogeneity without ACE2. Omicron_−ACE2_ has the broadest PC1 spread whereas Omicron_+ACE2_ shows more constrained PC1 (Figure 2b), indicating that ACE2 induces partial stabilization in Omicron, but less pronounced than in Delta. Figure 2c shows the change in the RBD-up position in open, full-open and extra-open conformational states, resulting an alteration of the dihedral angle of NTD-SD2-SD1-RBD and the RBD-core compactness. We observed Delta_+ACE2_ exhibits a peak at a distinct dihedral angle (~ −200°) whereas the other three systems show overlapping peaks around +180° (Figure S3a). Omicron_+ACE2_ shows a sharp peak, indicating a rigid and locked conformation. To investigate inter-RBD packing, we calculated the triangle area formed by residue TYR449 from each of the three protomers. Omicron with or without ACE2, and Delta in the absence of ACE2 exhibit a relatively compact RBD arrangement (Figure S3b). In contrast, only Delta shows a clear swelling of the RBD core upon ACE2 engagement, suggesting a variant-specific structural reorganization that is induced by receptor binding (Figure S3b).

**Figure 2.**
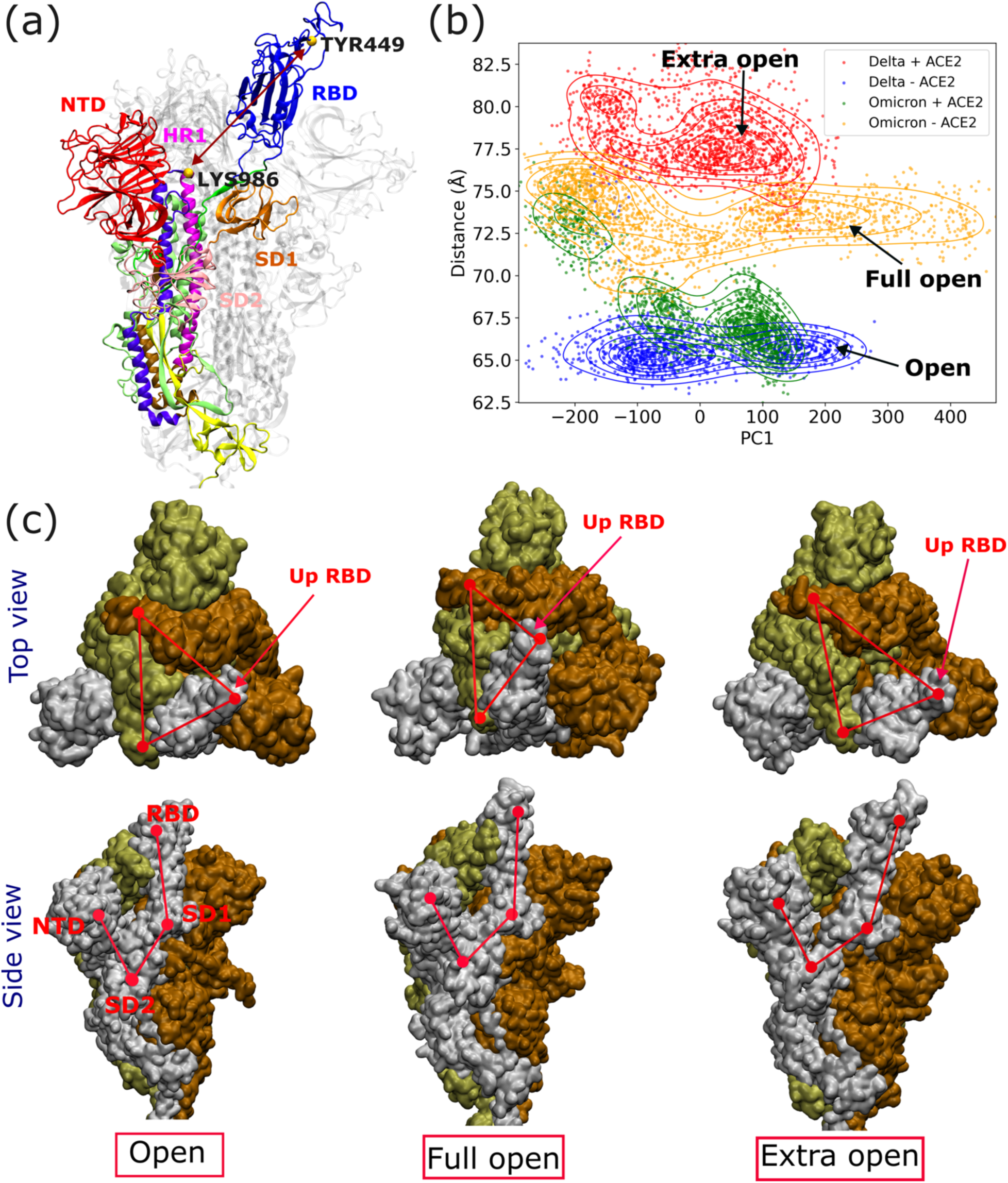
Structural definition and conformational distributions of the spike protein in Delta and Omicron variants. (a) Cα Distance between residue TYR449 and residue LYS986, reflecting the extent of RBD opening. (b) Principal component analysis projection onto the PC1 and TYR449-LYS986 distance for Delta and Omicron spike variants in the presence (+ACE2) and absence (-ACE2) of the receptor. (c) Representative conformations of the open, full open, and extra open states with corresponding triangular areas and dihedral angles.

Two-dimensional free energy landscapes were constructed by computing the joint probability distribution of the selected CVs, which were then transformed into a free energy surface using Boltzmann inversion. Figure 3 reveals distinct conformational preferences between Delta and Omicron upon ACE2 binding. In Delta, ACE2 promotes further opening of the RBD and stabilizes an expanded RBD core (Figure 3a, b). In contrast, Omicron exhibits multiple low-energy basins, including one near the fully open state (~75 Å) and another near the open state (~65 Å), indicating an ACE2-induced population shift toward a more compact RBD core (Figure 3c, d). These results suggest that Delta undergoes increased RBD opening in the presence of ACE2, transitioning from a “full open” to an “extra open” conformation. Conversely, Omicron displays the opposite trend, adopting a tighter open conformation upon ACE2 binding. Notably, the ACE2-driven population shifts differ between variants, consistent with prior reports that Delta samples more open conformations while Omicron exhibits tighter RBD-RBD interfacial packing and a more compact architecture.(9,24,71) In the following sections, we delineate the allosteric communication networks in Delta and Omicron, and correlate how pathway reconfiguration impacts the functional dynamics of the spike protein.

**Figure 3.**
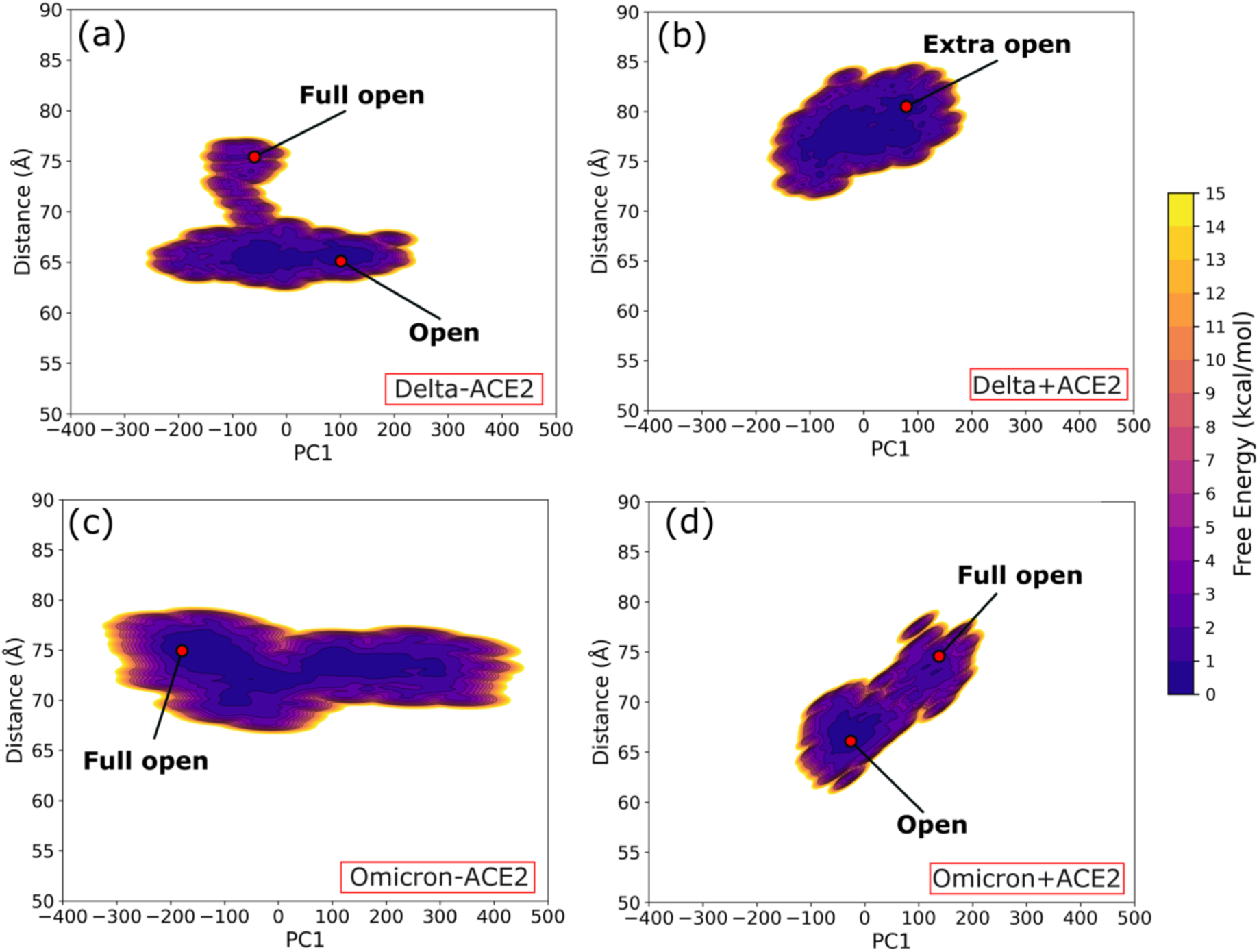
Free energy landscapes (FELs) of Delta and Omicron spike variants projected onto PC1 and the TYR449-LYS986 distance. Panels (a) and (b) show the FELs for Delta in the absence (-ACE2) and presence (+ACE2) of ACE2, respectively, while panels (c) and (d) correspond to Omicron without (-ACE2) and with ACE2 (+ACE2). The color scale represents free energy in kcal/mol.

### Reshaping the allosteric communication pathways in Delta and Omicron

To investigate and evaluate the allosteric communication networks in the Delta and Omicron spikes, we constructed cross-correlation matrices based on the positions of different residues in 3D space. A conventional method is the dynamic cross-correlation matrix (DCCM)(72), however, this method ignores correlated motions in orthogonal directions.(73) To overcome this limitation, we used an LMI-based cross-correlation matrix. LMI quantifies the strength of dynamic coupling between residue pairs more comprehensively.(26,74,75) A similar approach has been previously used to identify allosteric coupling during RBD opening in the SARS-CoV-2 wild-type variant.(26) In this work, we constructed residue-based networks using LMI calculations and compared the correlation heatmaps for Delta and Omicron in the presence and absence of ACE2.

The Spike, when unbound, exhibits widespread correlation between residues, indicating intrinsic interdomain communication and flexibility. Binding of ACE2 modulates this, channeling the landscape into more specific, and possibly functionally relevant, correlated motions. Figure S4 represents normalized LMI maps showing the residue-residue dynamic correlation in spike trimers. Delta exhibits a compartmentalization of correlations upon ACE2 binding, with smaller, more localized regions of strong communication. In contrast, Omicron displays enhanced inter-chain correlations with more continuous and brighter off-diagonal regions, signifying increased long-range communication and a dynamically connected trimer. Overall, these LMI heatmaps reveals that the ACE2 binding reshape the allosteric network differently in Delta and Omicron spikes.

To further probe the propagation of correlated motions, referred to as information flow, we constructed a protein connectivity network using a graph-theoretical approach(76), where each Cα atom is treated as a node and edges represent significant LMI-based correlations. We then computed betweenness centrality (BC), a graph-theoretical metric that quantifies how much a node mediates communication within the network(66,77) (see Methods). We calculated the difference in BC values {ΔBC = BC(+ACE2) - BC(-ACE2)} for both variants to identify key residues that gain or lose communication flow upon ACE2 binding (Figure 4a). We observed that ACE2 binding markedly enhances allosteric communication, involving more residues with higher BC values (BC > 0.2). We propose that this enhanced allosteric coupling helps channel information flow between different domains, including the NTD, Link1, Link2, SD1, SD2, Link3, FP, and CD, indicating that these regions play important roles in allosteric signaling within the spike protein (Figure 4a). Supporting this, hydrogen–deuterium exchange mass spectrometry (HDX-MS) experiments have shown increased deuterium exchange in the distant S1/S2 cleavage site and stalk region upon ACE2 binding in the S:ACE2 complex compared to free spike protein.(13) Another HDX-MS study by Chen *et al.* reports the receptor induced activation of spike proteins.(45) They observed a notable redistribution of dynamics across S1 and S2 upon ACE2 binding, reinforcing a receptor-triggered, network-wide response. Costello *et al.* reported that the population of the open state and the interconversion kinetics are strongly modulated by temperature, and are shifted by ACE2 binding, certain antibodies, and sequence variants, indicating a tunable energy landscape that links receptor/antibody engagement to global spike dynamics.(78)

**Figure 4.**
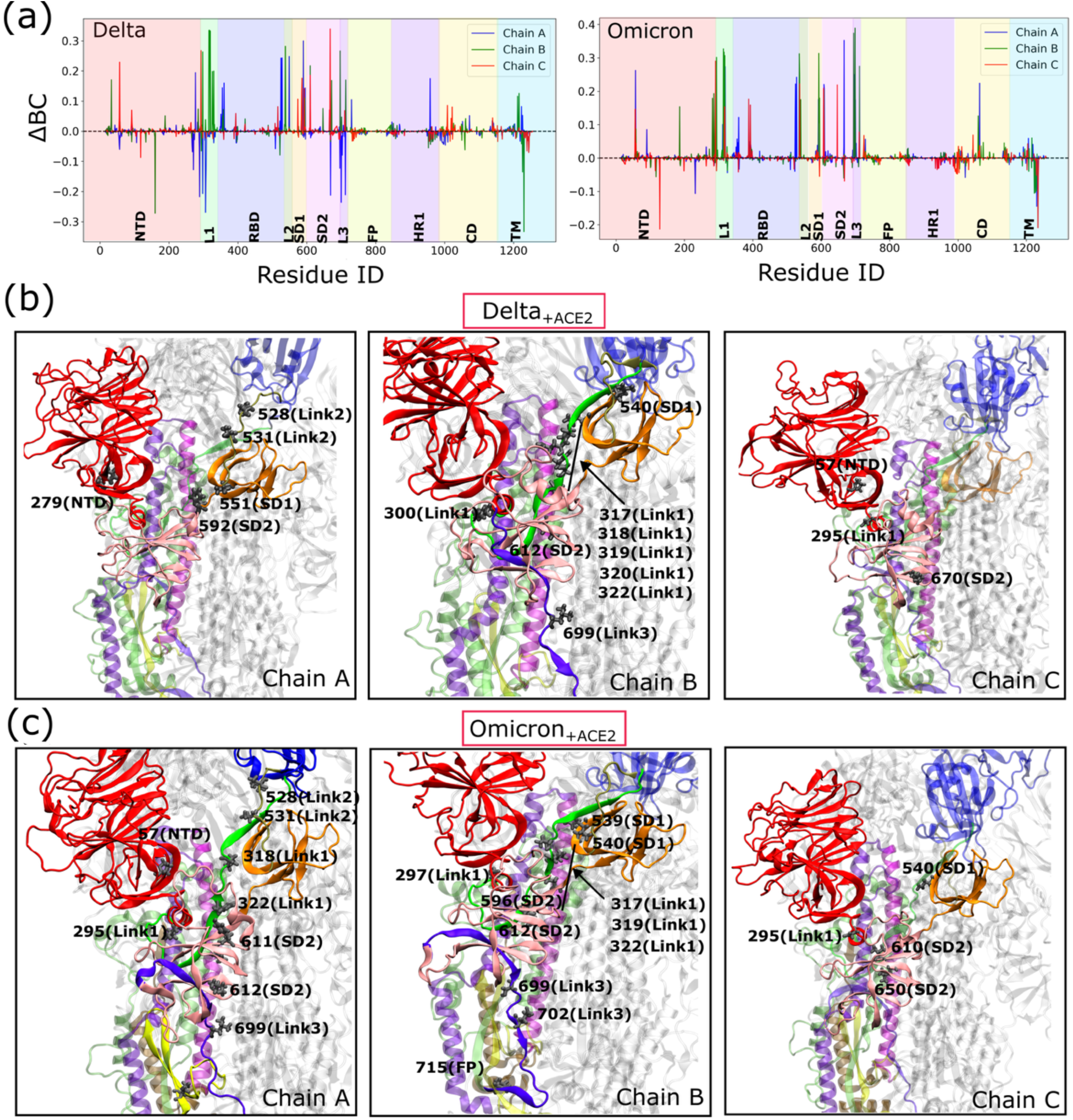
Modulation of allosteric communication in the SARS-CoV-2 spike protein. (a) Difference in betweenness centrality (ΔBC = BC[+ACE2] – BC[–ACE2]) per residue across chains A, B, and C upon ACE2 binding in Delta and Omicron. Residues with large positive or negative ΔBC values represent key nodes in ACE2-induced allosteric reorganization. Pannels (b) and (c) show the key resides with large changes in normalized BC values in Delta_+ACE2_ and Omicron_+ACE2_ systems, respectively.

To further identify the key residues that act as allosteric communication hubs, we selected residues with BC values greater than 0.2 (Figure S5). The key residues shared by both variants include residue 57 from the NTD; residues 295, 317, 318, 319, 320, and 322 from Link1; residues 528 and 531 from Link2; residues 539 and 540 from SD1; residues 611 and 612 from SD2; and residue 699 from Link3 (Figure 4b, c). Omicron uniquely gains long-range connectivity at residues 715 in the FP and 1066 in the CD. Consistent with our analysis, several experimental observations support receptor-induced long-range communications through the same structural corridors. In the NTD rim (proximal to residue 57 and the N61 glycan), glycan editing or enzymatic trimming at N17/N61/N165/N234 has been found to alter ACE2 binding and neutralization, remodel RBD opening, and shift antigenic exposure.(23,65,79,80) Cryo-EM(81,82) and smFRET(9) experiments on virion-displayed spike illustrate that perturbations at the NTD-RBD corridor (Link1: residues 295, 317-322) redistribute down to up state populations and modulate epitope accessibility, signifying that this band of residues participates directly in the opening hinge. At the Link2 (residues 528/531) and SD1 nodes (residues 539/540), variant and engineered substitutions in the same hinge neighborhood (for example, A570D) bias the RBD toward the up state,(83) while HDX-MS upon ACE2 binding exhibits enhanced protection around the RBD base/SD1,(84,85) together supporting the role of these residues in relaying receptor engagement to global conformational change. Additionally, in SD2 (residues 611/612, adjacent to D614), the well-characterized experimental studies demonstrate that D614G substitution reduces S1 shedding, increases spike stability, and shifts the population toward RBD-open ensembles,(86,87) strengthening the evidence that SD2 functions as a regulating node coupling S1 stability to RBD dynamics. In the Link3 corridor near the residue 699, changes at the neighboring furin loop (e.g., P681R in Delta) enhance S1/S2 cleavage and fusogenicity,(35,88) and ACE2 exposure primes downstream S2′ processing.(89) FP-proximal hub (residue 715) in Omicron aligns with studies that S2-side glycans and early-S2 segments (including the N717 region) tune fusion efficiency and trimer stability.(90,91) Additionally, CD/CH hubs (residue 1066) are supported by the prior structural and functional data showing that substitutions within the CH/CD bundle (e.g., N969K, L981F) reshape the S2 core, alter fusion kinetics, and stabilize prefusion architecture.(42,92,93)

To visualize the allosteric communication network between different structural subdomains of RBD-up and the RBD-down protomers, we constructed a domain-resolved heatmap by aggregating edge weights (LMI values) from the graph. For each edge connecting a source residue i and a sink residue j, the source and sink domains were identified based on residue indices, and the corresponding LMI value was added to the source-sink pair. Only edges with BC values greater than 0.2 were considered, thereby focusing the analysis on key residues likely to mediate significant information transfer.

Here, we present the source-sink heatmap for ACE2-bound systems only (Figure 5a, b), as ACE2-unbound systems exhibit negligible residues with BC > 0.2, indicating weaker coupling in the absence of the receptor. Notably, we observe that all three linkers (Link1, Link2, and Link3) show stronger coupling with the rest of the spike subdomains; however, the allosteric pathways undergo variant specific rewiring, modulating the strength of these sites across variants. Omicron exhibits a more pervasive communication network and gains long-range coupling between the S1 and CD domains, whereas in Delta, the network remains more localized within the S1 domain (Figure 5c, d).

**Figure 5.**
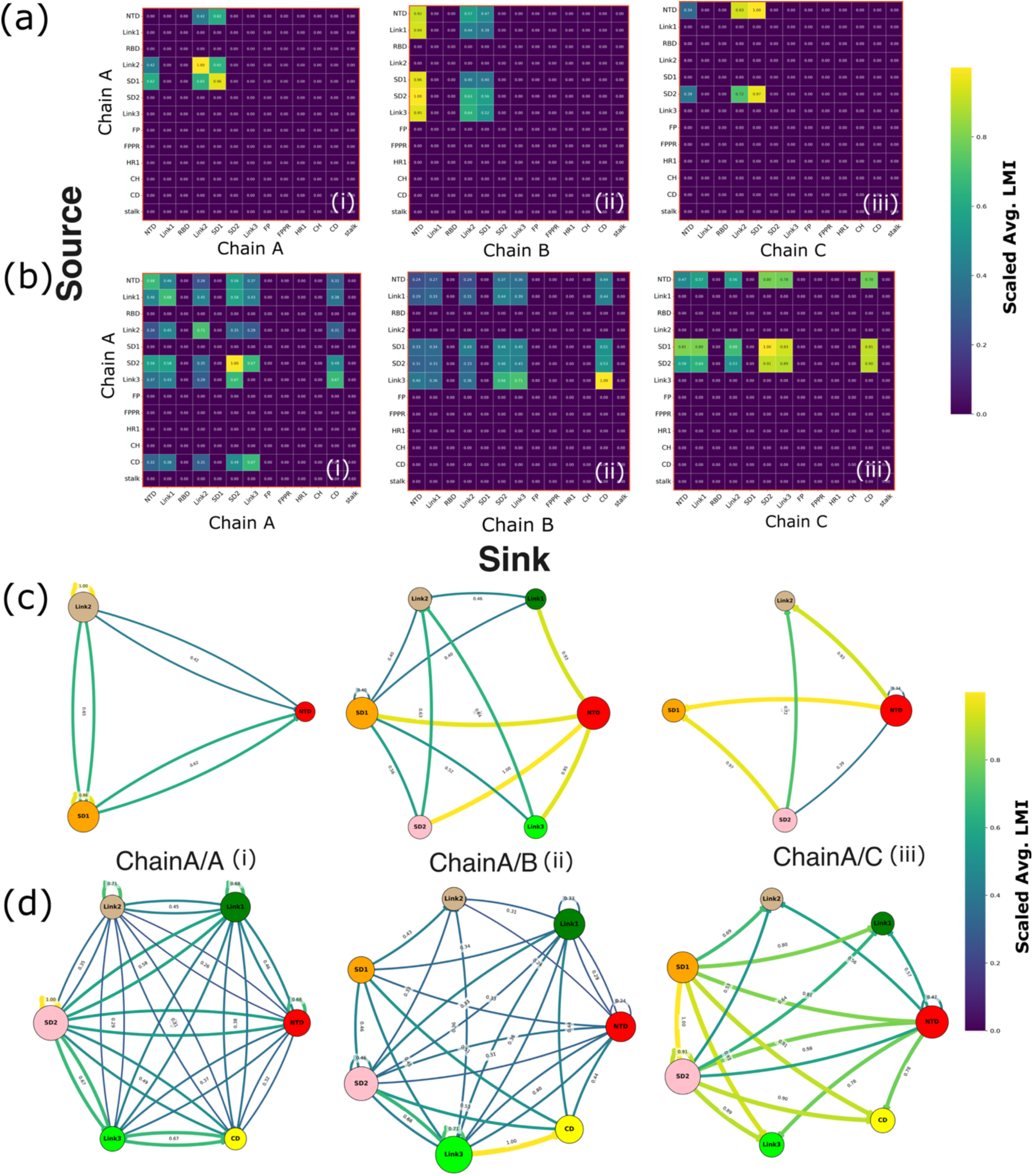
Source-sink allosteric communication analysis of the spike trimer. Scaled average LMI matrices showing interdomain coupling in the spike trimer for Delta+ACE2 (a) and Omicron+ACE2 (b), computed between defined structural domains. Panels (c) and (d) show the corresponding network representations, with nodes representing spike subdomains and edges indicating significant communication links. Panels (i)–(iii) indicate measurements for chain A/A, chain A/B, and chain A/C, respectively.

In Delta, intra-domain cross-talk within the RBD-up protomer (chain A) is primarily mediated through Link2 and SD1 (Figure 5a(i), c(i)), whereas Omicron gains connectivity in the SD2, Link3, and CD (Figure 5b(i), d(i)). Interestingly, the information flow from chain A to the neighboring chain B in Delta mostly passes through the NTD of chain B and the NTD, Link1, SD1, SD2, and Link3 of chain A (LMI > 0.8) (Figure 5a(ii), c(ii)). Although Omicron shows pervasive connectivity between chain A and chain B, the network appears weakened overall, except for strong interactions between Link3 of chain A and CD of chain B (LMI > 0.8) (Figure 5b(ii), d(ii)). In Delta, the NTD and SD2 of chain A exhibit stronger connectivity with Link2 and SD1 of chain C (Figure 5a(iii), c(iii)). In contrast, Omicron establishes connectivity between the SD1 and SD2 regions of chain A and the SD2, Link3, and CD regions of chain C (Figure 5b(iii), d(iii)). Overall, these results suggest that Delta exhibits a confined but stronger coupling via Link1 and Link2 within the S1 domains, whereas Omicron displays a more diffuse but longer-range network spanning S1 and S2 domains via Link3, indicating a rerouting of allosteric communication pathways. Several structural studies have shown that the extent of RBD opening has varied over viral evolution. Calvaresi *et al.* reported that early variants (Alpha, Beta, Delta) show greater opening upon ACE2 binding, whereas Omicron favors predominantly closed-like conformations, with SD1 acting as a lever that coordinates RBD, SD2, and FPPR across protomers.(84) Our results refine this picture: Link1 (NTD-RBD) modulates RBD opening, Link2 (RBD-SD1) governs S1 shedding, and Link3 (S1-S2 core) regulates priming/fusion. Upon ACE2 binding, Delta strengthens correlations through Link1/Link2, stabilizing an extra-open RBD state and promoting S1/S2 cleavage,(89) whereas Omicron gains connectivity through Link3, biasing the network toward fusion-priming control rather than extensive RBD opening(24).

Together, our analysis suggests that the three linker regions, Link1, Link2, and Link3, are critical mediators of the allosteric communication network extending from the ACE2 binding site to distant subdomains. Delta strengthening Link1 and Link2-mediated routes and Omicron gaining connectivity through Link3, which is therefore in line with experimental phenotypes: Delta’s greater fusogenicity and cleavage sensitivity(89) versus BA.1’s enhanced trimer stability, restrained opening, and remodeled antigenic surface(24,94).

### N856K and T547K mutations reshape the allosteric network in Omicron variant

Omicron has a significantly larger number of mutations on the spike surface than Delta (Figure 1c) and displays a higher surface density of positive charge compared with Delta (Figure S6). We propose that Omicron’s positively charged mutations play a significant role in reshaping the allosteric communication network. To investigate this, we identified salt-bridge residue pairs formed between the S1 domain of chain A the S1 and S2 domains of neighboring chains (Chain B and C). Additionally, we computed the residence time of these salt-bridge interactions to assess their stability and strength (Figure 6 and Figure S8). We observed the formation and disruption of several salt-bridge contacts across spike variants. In Omicron, the N856K mutation in chain B forms a salt bridge with ASP568 of chain A (Figure 6a). We also observed that the T547K mutation in Omicron establishes a stronger salt-bridge interaction with GLU748 upon ACE2 binding (Figure 6a). These two mutations contribute to significant conformational changes in the RBD, SD1, and SD2 regions of Omicron compared to Delta (Figure 6b). These salt-bridge interactions tightly anchor the tip of the S2 domain of chain B to the SD1 domain of chain A. We further note that another salt bridge between ARG567 and ASP979 appears to stabilize this interdomain contact. Such strong salt-bridge interactions between SD1 and the S2 tip hinder further RBD opening, resulting in a more compact RBD core in Omicron. We propose that the increased rigidity of the RBD core in Omicron reduces S1 domain flexibility, leading to a more pervasive but weaker allosteric network. Simultaneously, this rigidity maintains the S1-S2 interface, as reflected by a higher number of contacts between S1 and S2 domains (Figure S7), ultimately facilitating long-range coupling with the FP and CD domains.

**Figure 6.**
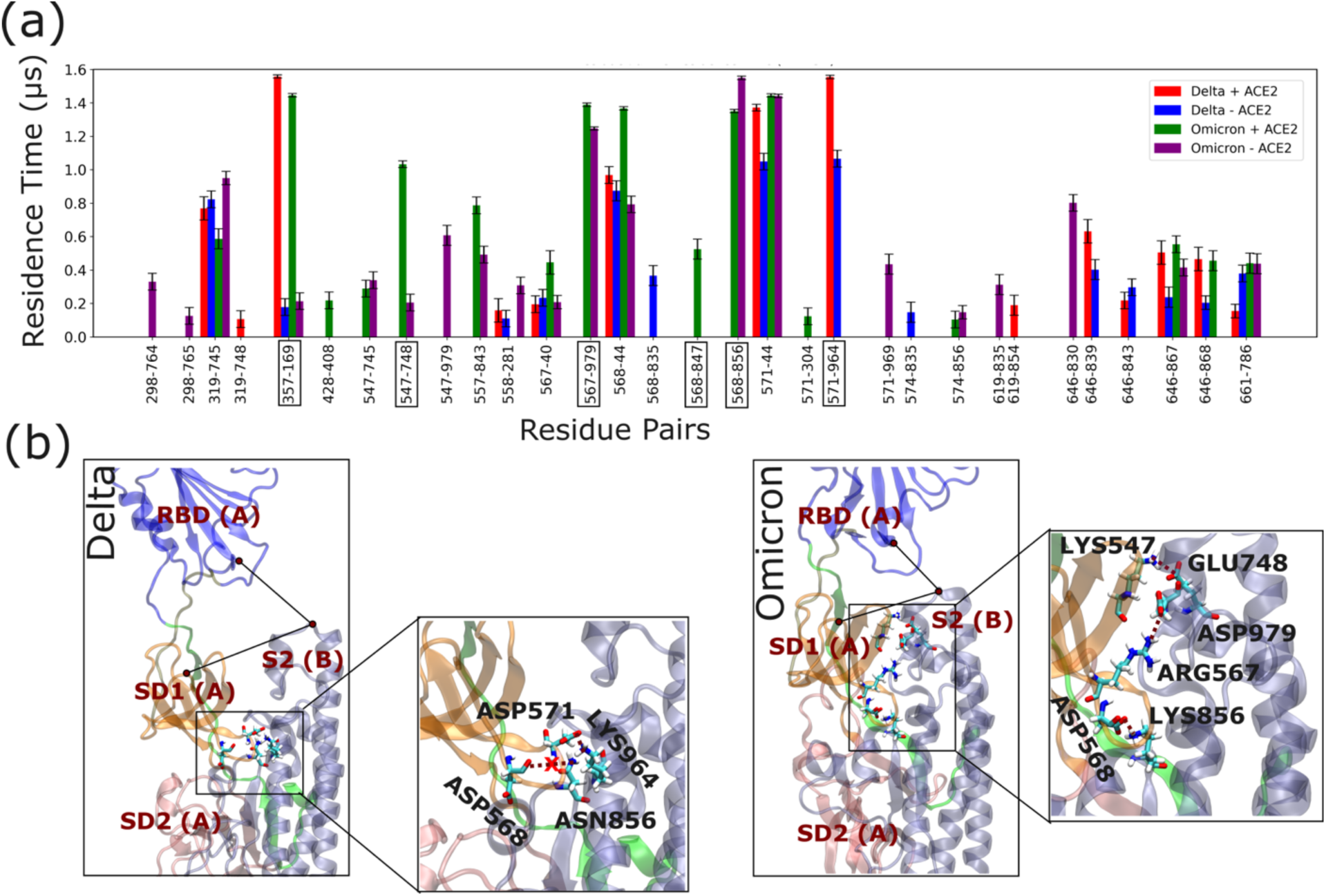
Variant-specific inter-protomer salt-bridge interactions and their structural localization in Delta and Omicron spike proteins. (a) Bar plot showing the residence times (μs) of inter-protomer (between chain A and chain B) salt-bridge residue pairs in Delta and Omicron spike variants in both ACE2-bound (+ACE2) and unbound (-ACE2) states. Here we considered the S1 domain of chain A and the S1 and S2 domains of chain B. The black box highlights key interactions that appear important for reshaping allosteric networks. (b) Representative structural snapshots highlighting key inter-protomer interactions: ASP571-LYS964 in Delta; ASP568-LYS856, LYS547-GLU748, and ARG567-ASP979 in Omicron.

Delta, on the other hand, forms a strong salt-bridge interaction between ASP571 (chain A) and LYS964 (chain B) (Figure 6a). This interaction positions the SD1 domain close to the HR1 helix but notably keeps it distant from the S2 tip, thereby displacing the SD1 domain of the RBD-up protomer away from the S2 region. This conformational rearrangement facilitates further opening of the RBD (Figure 6b). The increased openness of the RBD core enhances the flexibility of the S1 domain, which we propose contributes to a more focused and strengthened allosteric communication network. However, we also suggest that the extra-open RBD conformation leads to a significant detachment between the S1 and S2 domains (Figure S7), resulting in a loss of long-range connectivity.

Figure S8 shows the salt-bridge contacts between chain A and chain C. We observe that the S1 domain of chain A interacts primarily with the S1 domain of chain C. Delta_−ACE2_, Omicron_−ACE2_, and Omicron_+ACE2_ systems exhibit a greater number of RBD-RBD salt-bridge contacts compared to the Delta_+ACE2_ system. We propose that the more compact, less open RBD core in these systems facilitates the formation of RBD-RBD contacts. In contrast, the Delta_+ACE2_ system displays fewer RBD-RBD and more NTD-RBD salt-bridge interactions, which likely contribute to the stabilization of the extra-open RBD core conformation.

Additionally, we observe significant structural rearrangements in the 630-loop region (residues 620-640) of both Delta and Omicron variants (Figure S9). Structural and biochemical studies by Chen and co-workers have shown that the D614G mutation induces substantial structural and dynamic changes in 630-loop.(32) Furthermore, Dokainish *et al.* proposed a correlation between RBD opening and 630-loop movement.(95) In our analysis, the loop shifts toward the SD1 and SD2 domains and remains more connected to the S1 core in Omicron,, whereas in Delta, it moves away from SD1 and SD2 and becomes more disconnected from S1 (Figure S9). We identified an intra-loop salt-bridge interaction between ARG634 and ASP627 in Omicron, while in Delta, ARG634 forms a contact with ASP293 in the NTD. These observations are in line with the recent studies demonstrated that the 630-loop is directly involved in the NTD-to-RBD allosteric network in Omicron, while in Delta, it remains largely disconnected during the RBD’s closed-to-open transition.(27) We believe that the highly connected 630-loop in Omicron facilitates coupling between the S1 and S2 cores, whereas in Delta, its disconnection contributes to the loss of long-range S1-S2 communication.

## Conclusions

We have performed extensive atomistic MD simulations of the Delta and Omicron spike proteins in both ACE2-bound and unbound states to investigate their allosteric communication pathways. We observed that ACE2 binding stabilizes an additional open state of the RBD in Delta, whereas Omicron exhibits a population shift toward a less open RBD state upon ACE2 binding. Based on LMI calculations and graph theory-based analysis, we have also unraveled long-range allosteric communication networks within the spike proteins. Using betweenness centrality analysis, we identified residues critical for mediating maximal information flow across the spike.

With the RBD open, ACE2 engagement primes the spike for membrane fusion. The post fusion transition entails S1 release and refolding of S2, with HR1 and CH merging from short antiparallel helices into a continuous, elongated stem helix. Our results suggest that the distribution of ACE2 binding energy across S1 and S2 subunit is variant specific. Delta favors open conformations and channels its binding energy to open RBD protomers. Omicron, by comparison, favors the closed like state, and it allocates most of its binding energy toward S2, positioning the trimer for higher stability and immune escape. Mechanistically, three key linkers, Link1 (between NTD and RBD), Link2 (between RBD and SD1), and Link3 (between SD2 and FP) mediate the majority of allosteric communication between spike subdomains. Delta strengthens Link1 and Link2, producing confined, robust connectivity within S1 but diminished long-range coupling to the S2 core (Figure 7a), while Omicron redirects the network by reinforcing Link3, yielding a broader yet weaker S1 network alongside long-range connectivity with the CD domains (Figure 7b).

**Figure 7.**
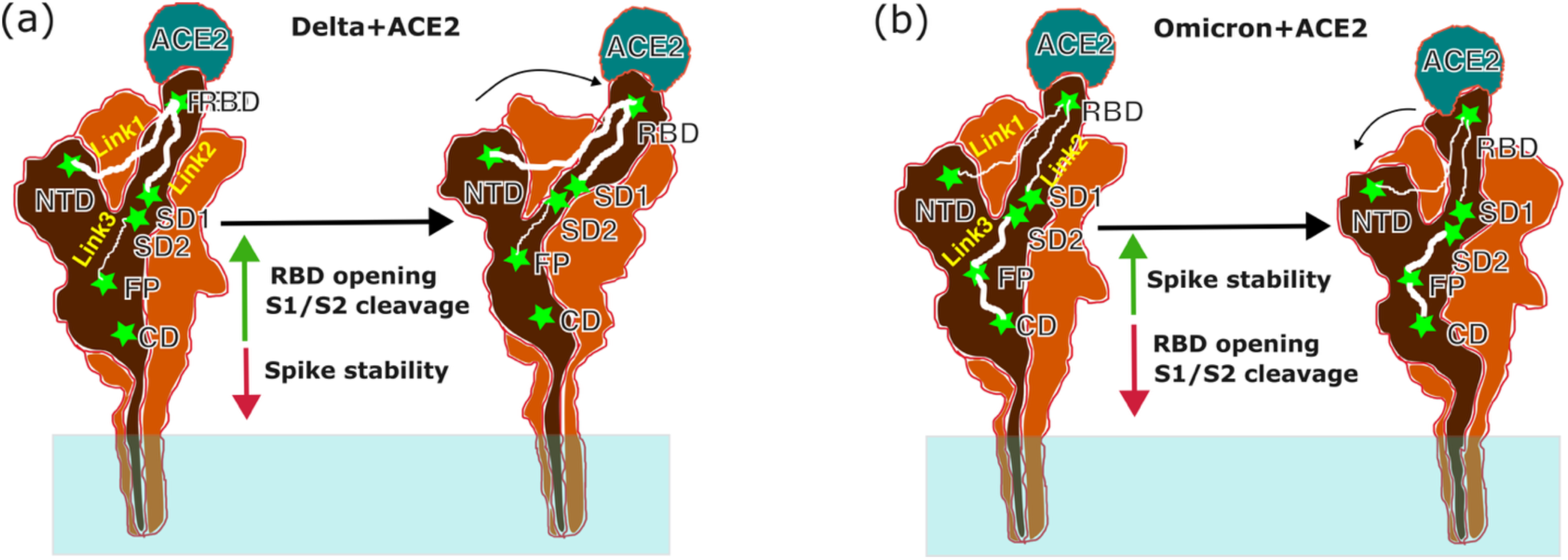
Schematic of the allosteric network in Delta and Omicron spike proteins. (a) Delta shows stronger connectivity within S1; Link1 and Link2 mediate communication, promoting greater RBD opening and increased S1/S2 cleavage. (b) Omicron shows long range connectivity from the RBD to the S2 core mediated via Link3, resulting in restrained RBD opening and enhanced trimer stability.

We further suggest that the N856K and T547K mutations remodel the conformational landscape, thereby reconfiguring the allosteric communication pathways in Omicron. Specifically, salt bridge interactions between ASP568-LYS856 and LYS547-GLU748 stabilize the interface between the tip of the S2 region and the SD1 domain of the RBD-up protomer, preventing further opening of the RBD. We also suggest that the compact RBD core in Omicron enables a more pervasive allosteric network, despite a reduction in overall connectivity strength. This configuration also preserves the S1-S2 interface, facilitating long-range coupling with distal regions such as the FP and CD. In contrast, Delta forms a strong salt bridge between ASP571 and LYS964, anchoring the SD1 domain near the HR1 helix but away from the S2 tip, thereby promoting an “extra-open” RBD conformation. We propose that this open RBD core enhances S1 domain flexibility and contributes to a more localized strengthening of the allosteric network. However, this conformational state also leads to a significant detachment between S1 and S2, resulting in diminished long-range allosteric connectivity.

Characterizing the fitness landscape of Spike mutations through thermodynamic analysis and graph theory-based network modeling can provide deeper insights into viral transmissibility and evolution in humans. The older variant Delta was favored for open spikes that aided receptor binding before RBD-targeted antibodies were prevalent. In contrast, Omicron likely reflects selection for less open spikes that shield RBD epitopes from widespread antibodies while retaining infectivity, potentially by lowering the energy barrier to fusion. Our study may open avenues for therapeutic intervention by identifying regions outside the RBD as potential inhibitor targets. By uncovering allosteric sites and mutation-driven conformational changes, our findings may help to lay the groundwork for designing antiviral strategies with enhanced resilience against emerging variants.

## Supporting information

Supporting Information Mutation and ACE2-induced Allosteric Network Rewiring in Delta and Omicron SARS-CoV-2 Spike Proteins

## Acknowledgments

This work was supported by the National Institutes of Health (NIH) grant R01AI169896. Computational resources were provided by the Frontera Supercomputer at the Texas Advanced Computer Center (TACC). Funded by the National Science Foundation (NSF grant OAC-1818253). Some initial equilibrations of the systems were performed on Midway2 at the Research Computing Center (RCC) at the University of Chicago.

## Author Contributions

M.D. and G.A.V. designed, M.D. performed, and both analyzed molecular dynamics simulations. M.D. and G.A.V. wrote the paper.

## Declaration of Interests

The authors declare no competing interests.

## Supporting Material

Document S1. Figures S1–S9

## Notes

### Competing Interest Statement

The authors have declared no competing interest.

