## Supporting Information Mutation and ACE2-induced Allosteric Network Rewiring in Delta and Omicron SARS-CoV-2 Spike Proteins for "Mutation and ACE2-induced Allosteric Network Rewiring in Delta and Omicron SARS-CoV-2 Spike Proteins"

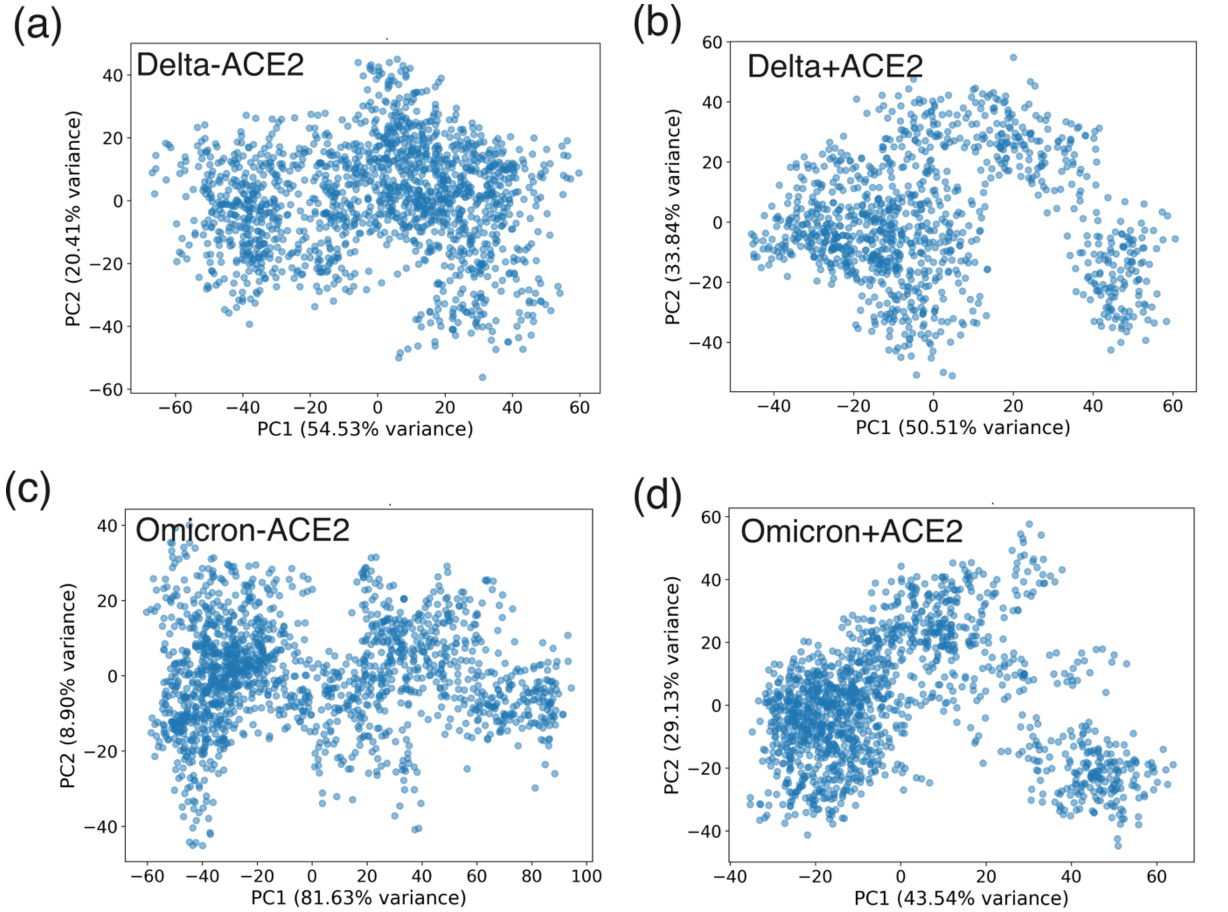


**Figure S1.** Principal component analysis of the spike protein dynamics of Delta and Omicron BA.1 variant in presence and absence of ACE2. Projection of first two principal components (PC1 vs. PC2) of the S1 domains of Delta-ACE2 (a), Delta+ACE2 (b), Omicron-ACE2 (c), Omicron+ ACE2 (d).

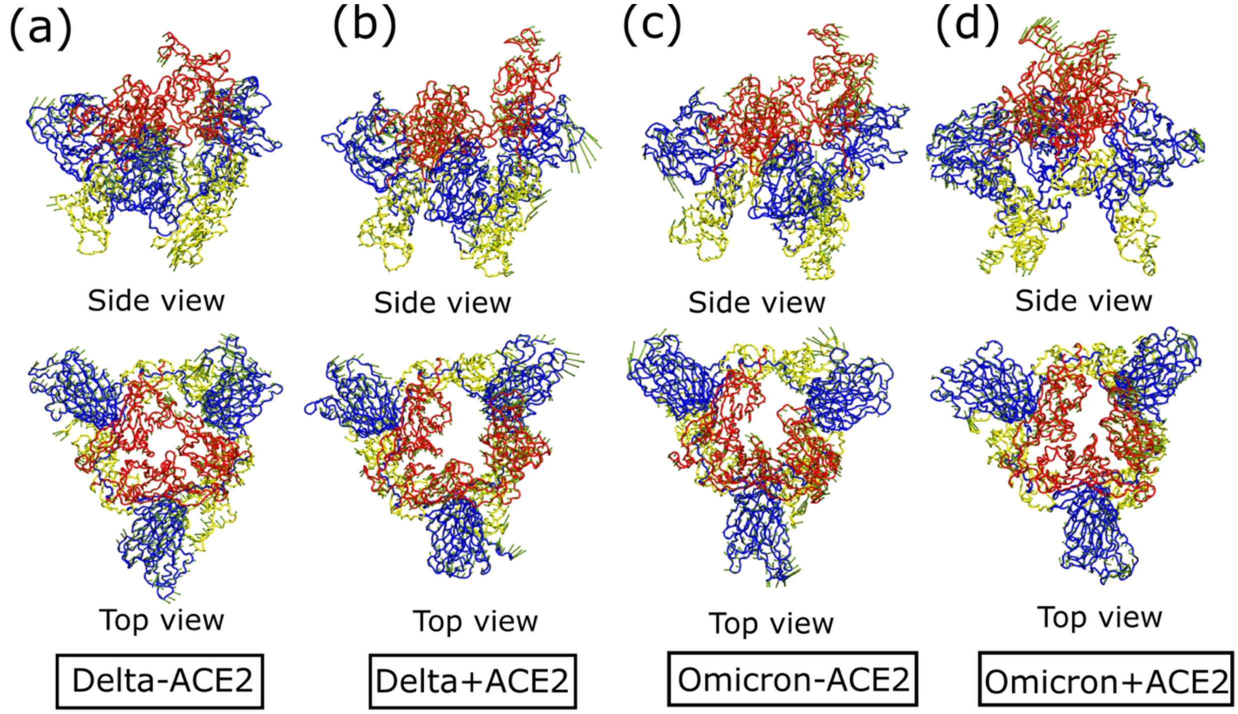


**Figure S2**. Representation of PC1 movements of S1 domain of spike variants. The red, blue and yellow colors represent NTD, RBD and SD (SD1/2) domains of Delta-ACE2 (a), Delta+ACE2 (b), Omicron-ACE2 (c), Omicron+ACE2 (d), respectively.

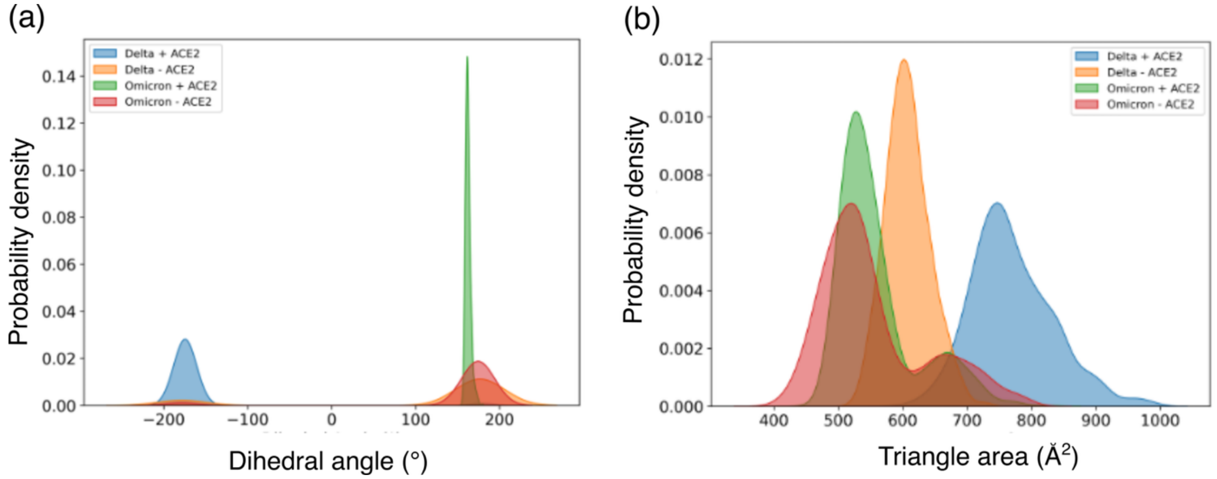


**Figure S3.** Distribution plots of dihedral angle and triangle area of the spike S1 domains in Delta and Omicron variants. (a) Probability density distribution of the dihedral angle calculated using the center of mass of the NTD, SD2, SD1, and RBD domains, comparing Delta and Omicron in the presence and absence of ACE2. (b) Probability density distribution of the triangle area formed by residue TYR449 from each of the three protomers, highlighting differences in inter-RBD spatial arrangement across variants and ACE2 binding states.

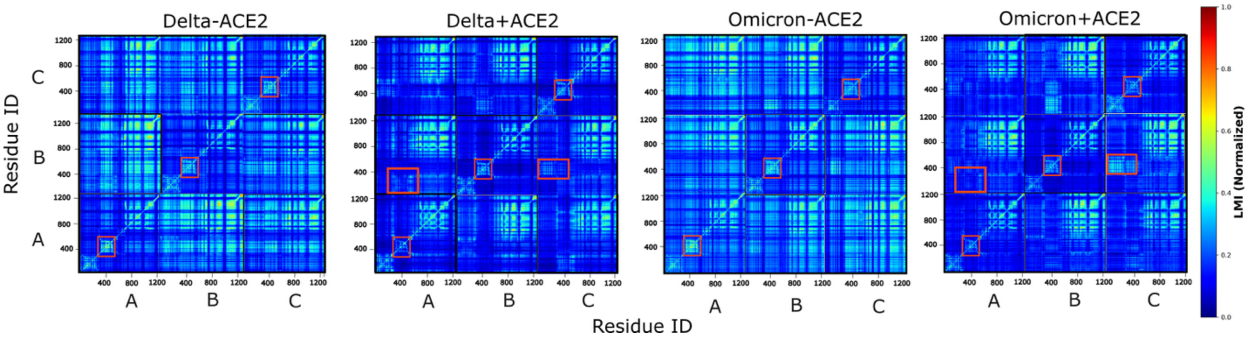


**Figure S4.** Normalized LMI maps showing residue-residue dynamic correlations in the spike trimer of Delta and Omicron variants. Heatmaps represent normalized LMI values between Cα atoms of residues in Delta and Omicron spike proteins in both ACE2-unbound and ACE2-bound states. The red boxes highlight key regions where gain or loss of dynamic correlation is observed across different simulated systems.

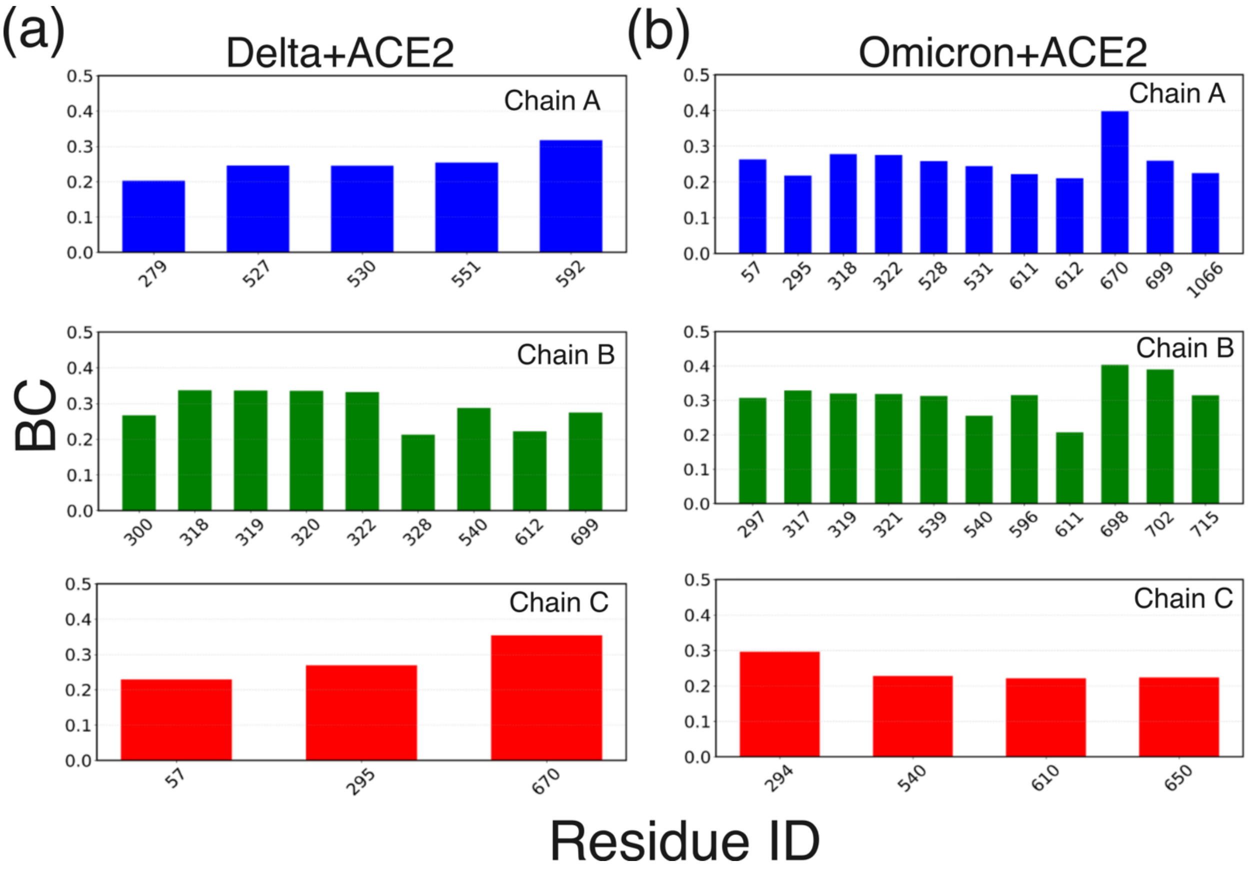


**Figure S5.** BC values of key residues in the spike trimer of Delta and Omicron variants in the ACE2-bound state. Pannels (a) and (b) shows the key resides in Delta$_{+ACE2}$ and Omicron$_{+ACE2}$ systems, respectively.

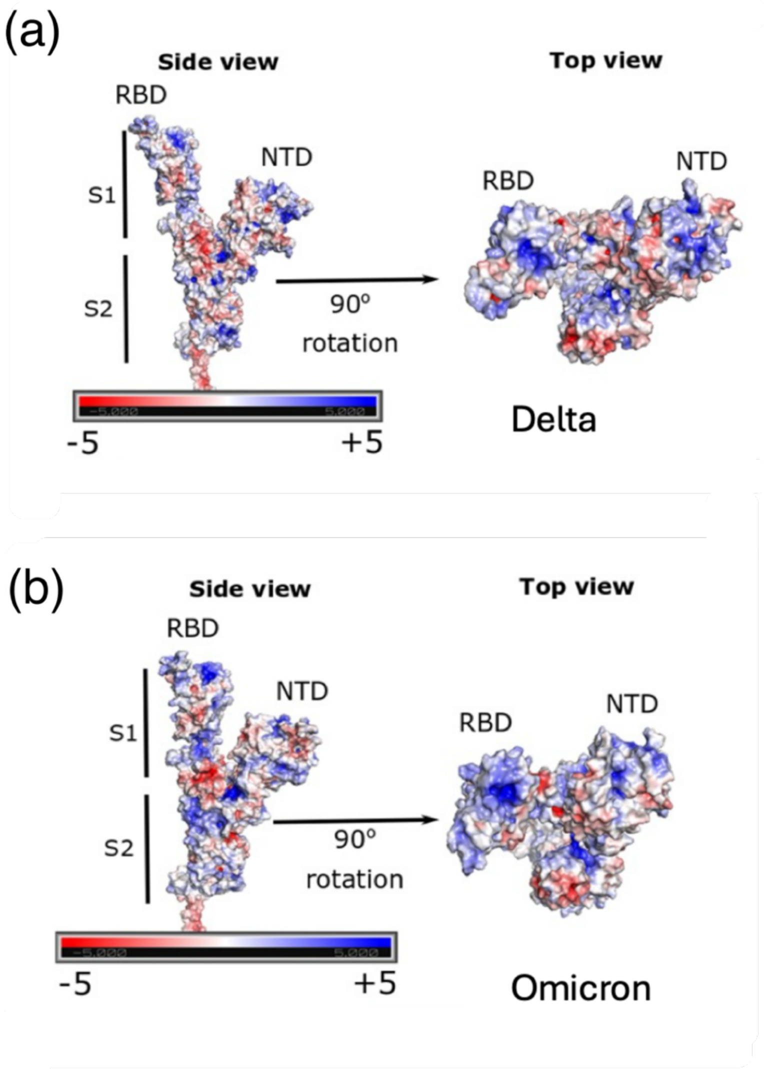


**Figure S6.** Electrostatic surface representations of the SARS-CoV-2 spike protein. Panels (A) and (B) show the electrostatic surface plot of the spike protein for the Delta and Omicron variants, respectively.

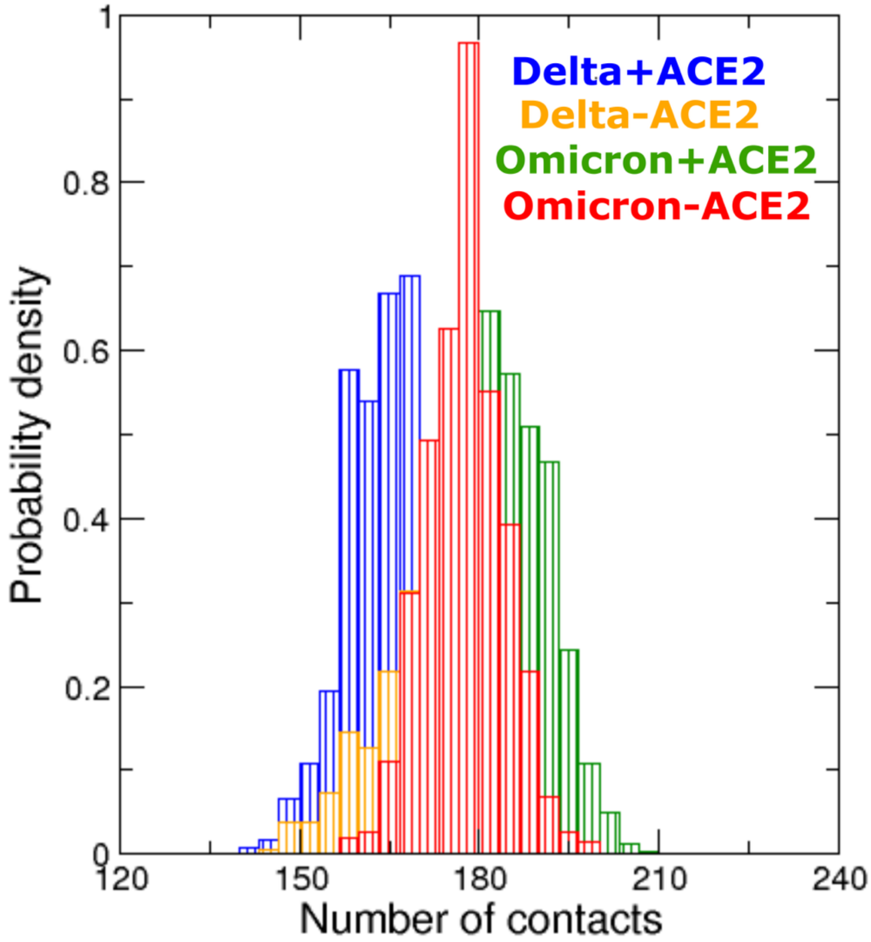


**Figure S7.** Probability density distribution of the number of contacts between the S1 and S2 domains of spike proteins in the presence and absence of ACE2. The distribution is calculated using Cα atoms with a cutoff distance of 0.6 nm to define contacts.

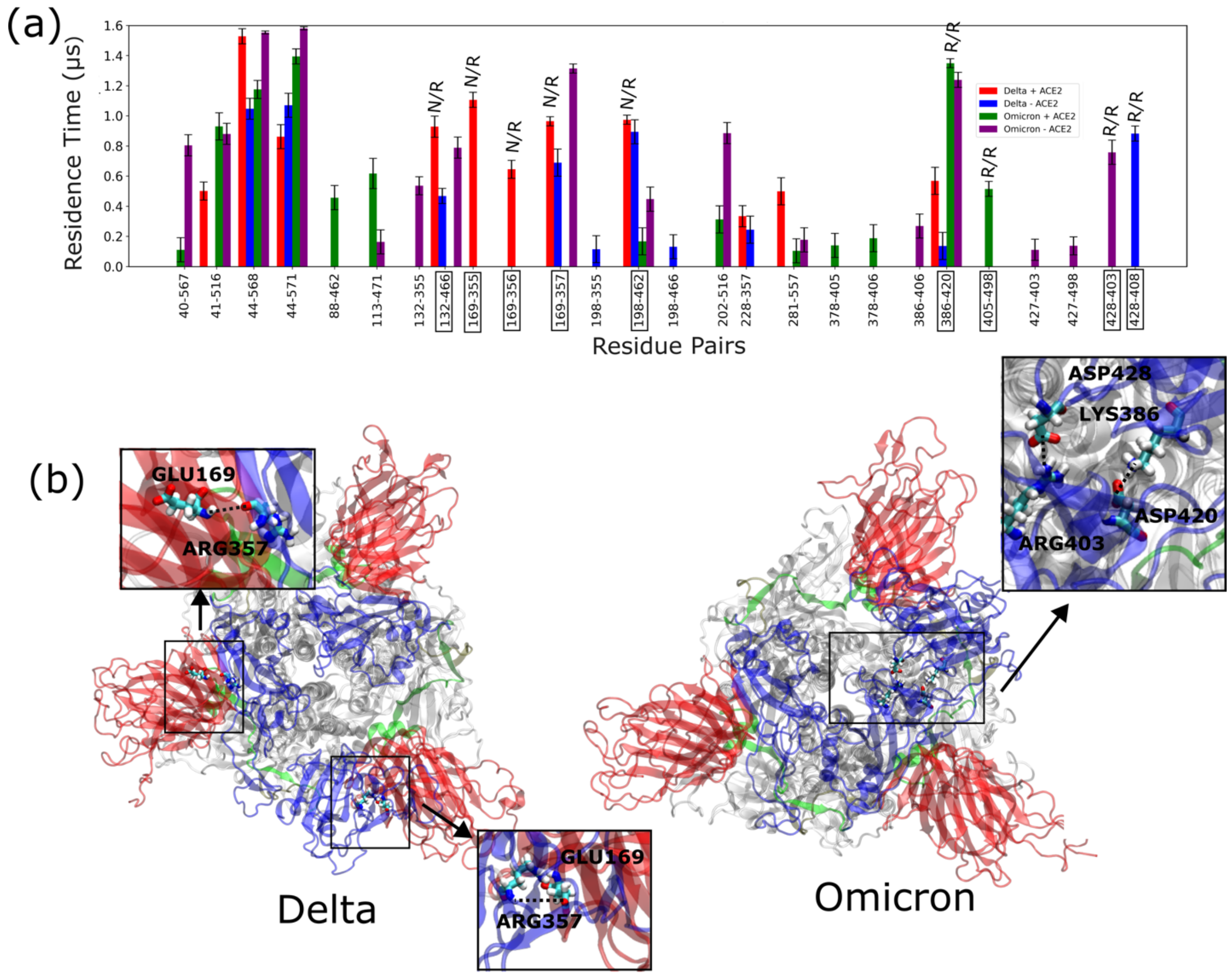


**Figure S8.** Variant-specific inter-protomer salt-bridge interactions and their structural localization in Delta and Omicron spike proteins. (a) Bar plot showing the residence times (μs) of inter-protomer salt-bridge interactions between chain A and chain C in Delta and Omicron spike variants, under both ACE2-bound (+ACE2) and unbound (-ACE2) conditions. We considered the S1 domain of chain A and the S1 and S2 domains of chain C. Key interactions are highlighted with black boxes. N/R and R/R represent NTD/RBD and RBD/RBD contacts, respectively. Notably, Delta shows more N/R contacts, whereas Omicron shows more R/R contacts. (b) Representative structural snapshots illustrating the spatial localization of key inter protomer salt bridge interactions identified in panel (a).

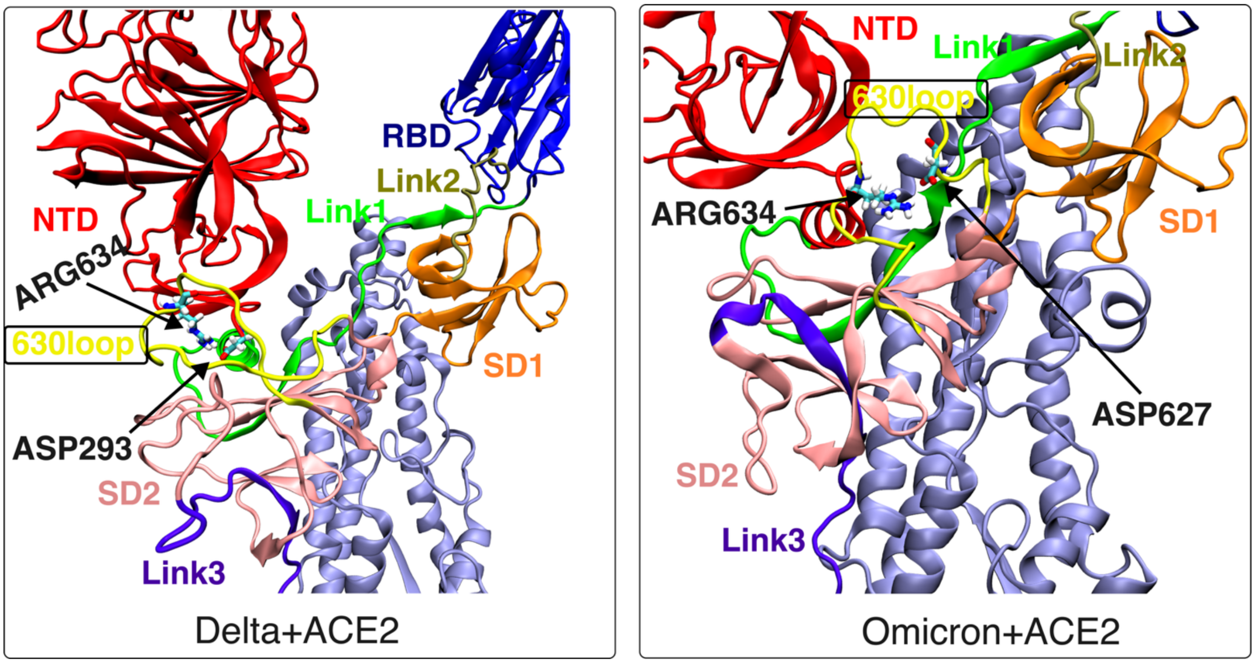


**Figure S9.** Orientation of the 630-loop in Delta and Omicron spike proteins in the ACE2-bound state. This figure highlights structural details of the 630-loop region, comparing its orientation in the Delta and Omicron variants upon ACE2 binding.
